# Cerebellar white matter development is regulated by fractalkine-dependent microglia phagocytosis of oligodendrocyte progenitor cells

**DOI:** 10.1101/2024.11.15.620441

**Authors:** McKenzie K. Chappell, John Shelestak, Muhammad Irfan, Eric Shelestak, Ashley D. Nemes-Baran, Gabrielle M. Mey, Tara M. DeSilva

## Abstract

Complex neurodevelopmental disorders involve motor as well as cognitive dysfunction and these impairments are associated with both cerebral and cerebellar maturity. A network of connections between these two brain regions is proposed to underlie neurodevelopmental impairments. The cerebellar gray matter has a protracted developmental timeline compared to the cerebral cortex, however, making the association of these relay pathways unclear for neurodevelopmental disabilities. We show that a population of amoeboid microglia infiltrate the cerebellar white matter through the fourth ventricular zone during early postnatal development. This infiltration is synchronized with the emergence of amoeboid microglia in the ventricular zone of the lateral ventricles and appearance in cerebral white matter. Amoeboid microglia phagocytosed oligodendrocyte progenitor cells (OPCs) in the cerebellar white matter during a restricted early postnatal time window before transitioning to a ramified morphology. Modulating fractalkine receptor signaling, shown to be involved in microglial pruning of synapses, significantly reduced microglial engulfment of OPCs resulting in increased numbers of OLs and altered myelin formation. Variants in the fractalkine receptor are associated with neurodevelopmental disorders including schizophrenia and autism where myelin perturbations have been documented. Overall, these data support that white matter refinement by amoeboid microglia is coordinated in both cerebral and cerebellar development with important implications for altered circuit function in neurodevelopmental disabilities.

**One sentence summary:** Microglia engulf oligodendrocyte progenitors in the developing cerebellum

## Introduction

The cerebellum plays a significant role in balance and coordinated movement, however, more recently it is receiving attention for its involvement in cognitive function such as modulation of attention, language, and social behavior^1,2^. The cerebellum receives input from the inner ear^3,4^, spinal cord^5–7^, and cerebral cortex^8–10^. One major pathway of communication between the cerebral and cerebellar cortices is the cortico-ponto-cerebellar pathway. Information from the cerebral cortex is transmitted via axons to the pontine nuclei^11,12^ and relayed to mossy fibers into the cerebellar cortex^13,14^. After integrating this information, Purkinje cells, whose axons traverse the cerebellar white matter to deep cerebellar nuclei^15^, transmit this information to different areas of the CNS. In fact, Purkinje cells are the only efferent system for the cerebellum and comprise the predominant axons in the cerebellar white matter and to a lesser extent mossy fibers.

Myelin is not only essential for action potential propagation, but also provides metabolic support to axons, supporting its role in neural function^16,17^. Proper myelination is also important in the development and formation of Purkinje axon plexuses in the granule cell layer^18^ and has been shown to have a retrograde inhibitory effect on the expression of growth-associated genes in Purkinje cells^19^. Severe myelin deficiency in Long Evans Shaker rats exhibited hyperexcitability of neurons in the deep cerebellar nuclei^20^. The role of Purkinje neurons as the sole output for the cerebellum and the importance of myelin in regulating Purkinje cell function highlights the significance of understanding mechanisms that regulate cerebellar white matter development.

Microglia contribute in many ways to the formation of proper neural circuits during development including synaptic pruning^21–23^. Derived from RUNX1+ primitive macrophages in the yolk sac, microglia begin populating the CNS as early as embryonic day 10 in mice^24^. We previously published that a population of amoeboid microglia arise in the ventricular zone of the lateral ventricles around term birth^25^. Amoeboid microglia invade the corpus callosum during early postnatal (P) development and engulf viable oligodendrocyte progenitor cells (OPCs) at P7 before assuming a resting morphology at the onset of myelination (P11)^25^. Modulating fractalkine receptor signaling, shown to be involved in microglial pruning of synapses, significantly reduced microglial engulfment of OPCs resulting in altered myelin formation^25^. While these data suggested that ameboid microglia refine the OPC pool as a homeostatic process for proper OPC-to-axon ratio during a specific timeframe in white matter development, it remained unclear if this process of refinement occurred in other white matter regions of the brain.

Since amoeboid microglia were found in the lateral ventricular zone adjacent to the corpus callosum, we investigated the 4^th^ ventricular zone adjacent to the cerebellar white matter, which undergoes myelination at a similar developmental timepoint to the corpus callosum^26–28^. Indeed, a population of amoeboid microglia was detected in the 4^th^ ventricular zone and migrated into the cerebellar white matter concurrent with their appearance in the corpus callosum. Amoeboid microglia also engulfed viable OPCs in the cerebellar white matter during a restricted window of development before the onset of myelination. Fractalkine receptor (CX3CR1) KO mice, one of the most widely used models to study alterations in microglia pruning of synapses^29^ also reduced engulfment of OPCs leading to hypomyelination. The conservation of this behavior in microglia indicates a need to control OPC numbers to modulate myelin formation in white matter tracks across the brain and that CX3CR1 signaling is a key regulator of this process. Overall, these studies have important relevance to neurodevelopmental disorders that affect cerebral and cerebellar function and are associated with both myelin alterations^30–33^ and variants in the fractalkine receptor^34^.

## Results

### Postnatal infiltration of amoeboid microglia from ventricular zones is synchronized in cerebellar and cerebral white matter

Using a green fluorescent reporter to label microglia (CX3CR1:GFP), cells with an amoeboid morphology were found in the fourth ventricular region at postnatal day (P) 0 (birth) (Fig. 1 A-B). The trunk of arbor vitae region of the cerebellar white matter is adjacent to the fourth ventricular zone and amoeboid microglia infiltrated this region by P3 (Fig. 1 C-D). As development progressed, amoeboid microglia increased in number with the majority present in the cerebellar white matter at P7, however, the presence of amoeboid microglia infiltration in the overlying cerebellar cortex was relatively few (Fig. 1 E-F). After this timepoint, amoeboid microglia assumed a ramified morphology (Fig. 1 G-H). The timeframe for the appearance of amoeboid microglia in the fourth ventricle is consistent with their appearance in the lateral ventricles (Fig. 1, arrows) and their early postnatal invasion into the adjacent corpus callosum as we published^25^. Taken together, these data support that the synchronized timing of amoeboid microglial infiltration into the cerebral and cerebellar white matter is part of an important neurodevelopmental process specific for white matter tracts in the brain.

**Figure 1.**
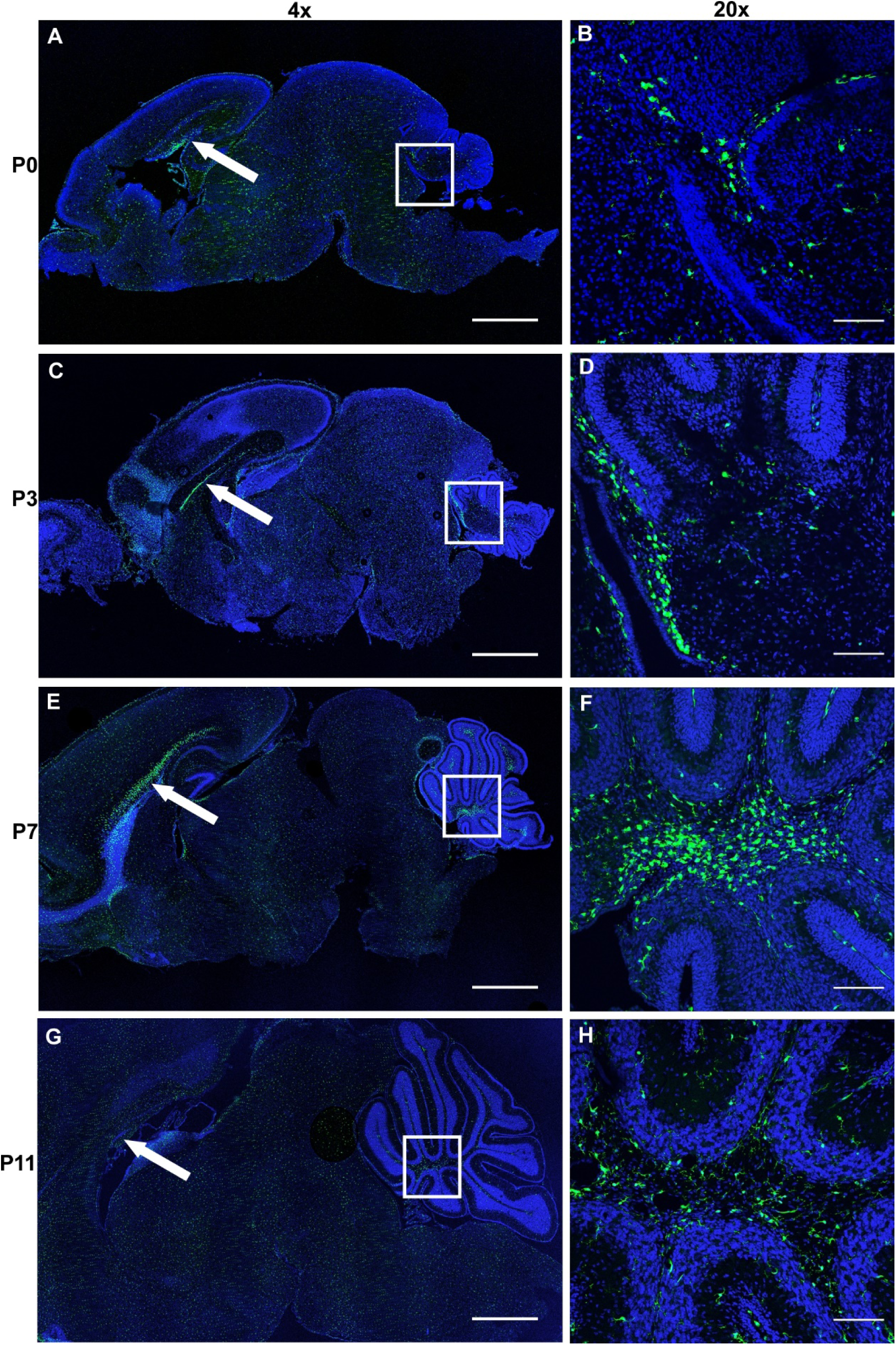
Postnatal infiltration of amoeboid microglia from ventricular zones is synchronized in cerebellar and cerebral white matter. Representative 4x images of sagittal brain sections from CX3CR1:GFP mice that fluorescently label microglia green, counterstained with bisbenzimide to visualize cellular nuclei at postnatal (P) day 0 (A), 3 (C), 7 (E) and 11(G). White boxes represent 20X magnification of the 4^th^ ventricular zone adjacent to the cerebellar white matter for each respective time point (B, D, F, and H). Arrows indicate the ventricular zone of the lateral ventricles below the corpus callosum. (A, B) Amoeboid microglia are present in the ventricular zone of the 4^th^ ventricle (white box) at the same time amoeboid microglia are observed in the ventricular zone of the lateral ventricles (white arrow). (C,D) Amoeboid microglia beginning to infiltrate the cerebellar white matter similar to invasion of amoeboid microglia into the corpus callosum. (E,F) At P7 amoeboid microglia predominantly occupy the cerebellar white matter, mirroring peak infiltration of amoeboid microglia into the corpus callosum. (G,H) By P11, microglia assume a resting morphology in the cerebellar white matter similar to the corpus callosum (white arrow). For each timepoint, *n* = 3 animals, 2 images at 4X and 2 images at 20X for each animal. Scale bars: 4x = 1000 µm; 20x = 100 µm.

### 3D Confocal reconstructions show amoeboid microglia phagocytosing OPCs within the cerebellar white matter at P7

The infiltration of amoeboid microglia into the cerebellar white matter during early postnatal development occurs during oligodendrogenesis^35^. To evaluate the interaction between amoeboid microglia with OPCs, CX3CR1:GFP mice were crossed with NG2:Tom mice to fluorescently label OPCs red. Confocal 3D reconstructions were collected during this restricted window of postnatal development when amoeboid microglia are present in the cerebellar white matter (P3-P11, Fig. 2). At P4, amoeboid microglia were found contacting OPCs with some evidence of engulfment (Fig. 2A). At P7, there was a dramatic increase in amoeboid microglia engulfment of OPCs (Fig. 2B), concurrent with amoeboid microglia engulfment of OPCs in the corpus callosum as we published^25^. 3D reconstruction of an amoeboid microglia (green, Fig. 2D) shows encapsulation of an OPC (red, Fig. 2E, video in supplemental data movie S1). Engulfment is also demonstrated in a single 0.3 µm plane from the z-stack (Fig. 2F-G). At the P11 time point, microglia exhibited a ramified morphology with little to no OPC engulfment (Fig. 2C), consistent with the onset of myelination. To evaluate if the observed engulfment of OPCs by amoeboid microglia at P7 in the cerebellar white matter was processed into the lysosome, immunofluorescence for CD68 a myeloid-specific lysosomal marker was used. Engulfment is defined as ≥15% of the NG2:Tom OPC volume encapsulated within microglia and contacting is defined as ≤15% of the NG2:Tom OPC internalized within microglia^25,36^. Confocal 3D reconstructions show amoeboid microglia (green, Fig. 3A) engulfing OPCs (red, Fig. 3B, merge in 2C) with CD68+ lysosome staining (pink, Fig. 3D) surrounding engulfed OPCs (merge in 3E), and colocalization with bisbenzimide (blue), a nuclear marker (Fig. 3F, merge all in 3G, video in supplemental data movie S2). The CD68+ lysosome occupied 65.19% ± 3.93 of microglia volume, while OPCs occupied 22.50% ± 3.84 (Fig. 3H). The majority of engulfed OPC volume was internalized within the CD68+ lysosome 68.09% ± 4.72 and was statistically increased compared to the engulfed OPC volume not within the lysosome 31.91% ± 4.72 (Fig. 3I). Therefore, these data provide evidence that OPCs internalized within microglia are being phagocytosed and targeted for degradation in the cerebellar white matter.

**Fig 2.**
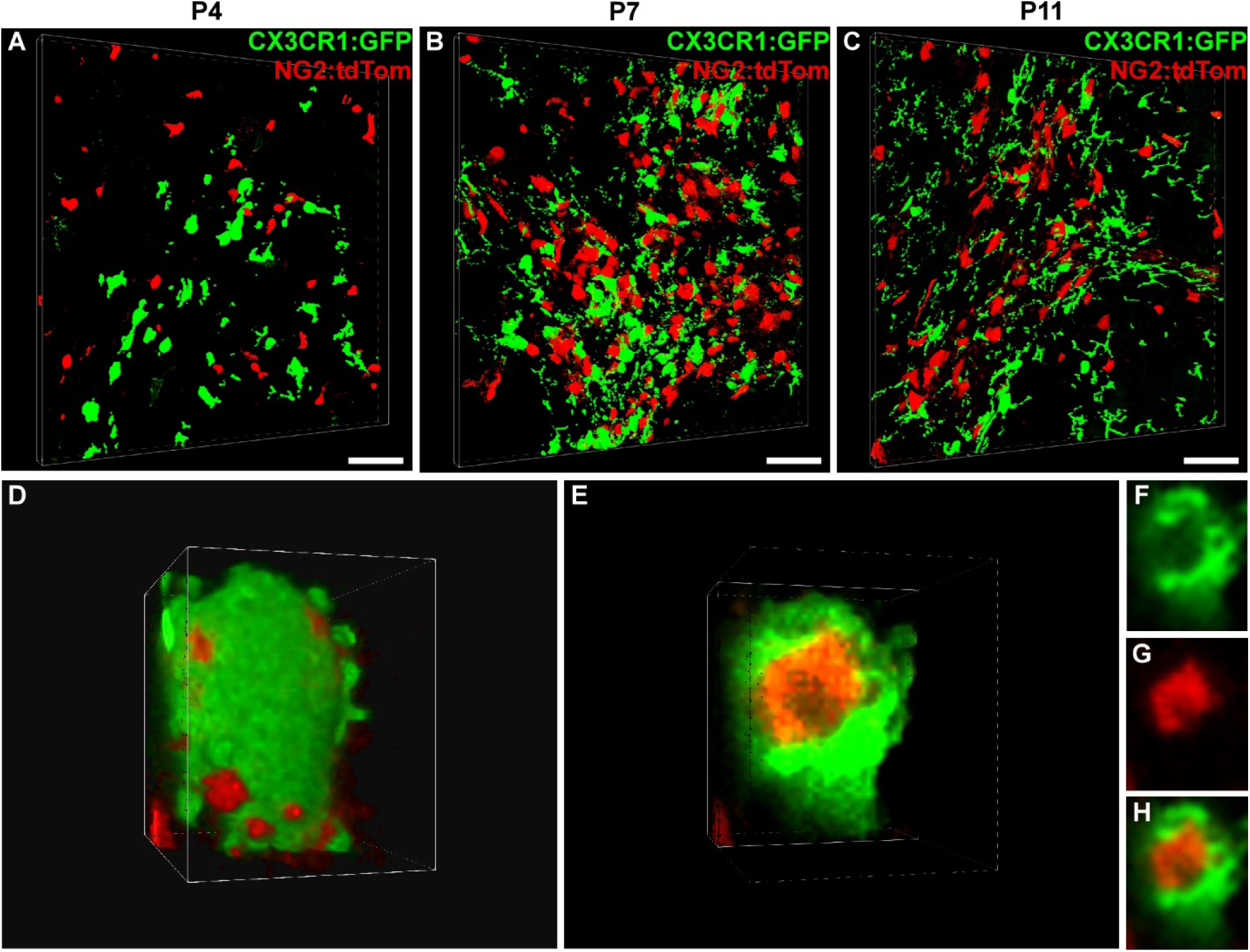
Microglial morphology changes across development and reflects peak engulfment in cerebellar white matter at postnatal day (P) 7. (A-C) Representative 3D confocal reconstructions from mice with green microglia (CX3CR1:GFP) and red oligodendrocyte progenitor cells (OPCs, NG2:Tom) during early cerebellar white matter development, scale bars = 50µm. (A) At P4, microglia interactions with OPCs are observed with limited engulfment. (B) At P7, amoeboid microglia are observed contacting and engulfing OPCs. (C) By P11 microglia exhibit a more ramified morphology with less evidence of engulfment. Amoeboid microglia from z-stack rendered in 3D (D). Cutaway into z-stack (D) shows an OPC encapsulated by a microglia (E, video in supplemental data movie S1). Single 0.3µm plane from z-stack showing microglia (F), OPC (G), and merge of both channels (H). *n* = 3 mice for each timepoint with three z-stacks taken per mouse.

**Figure 3.**
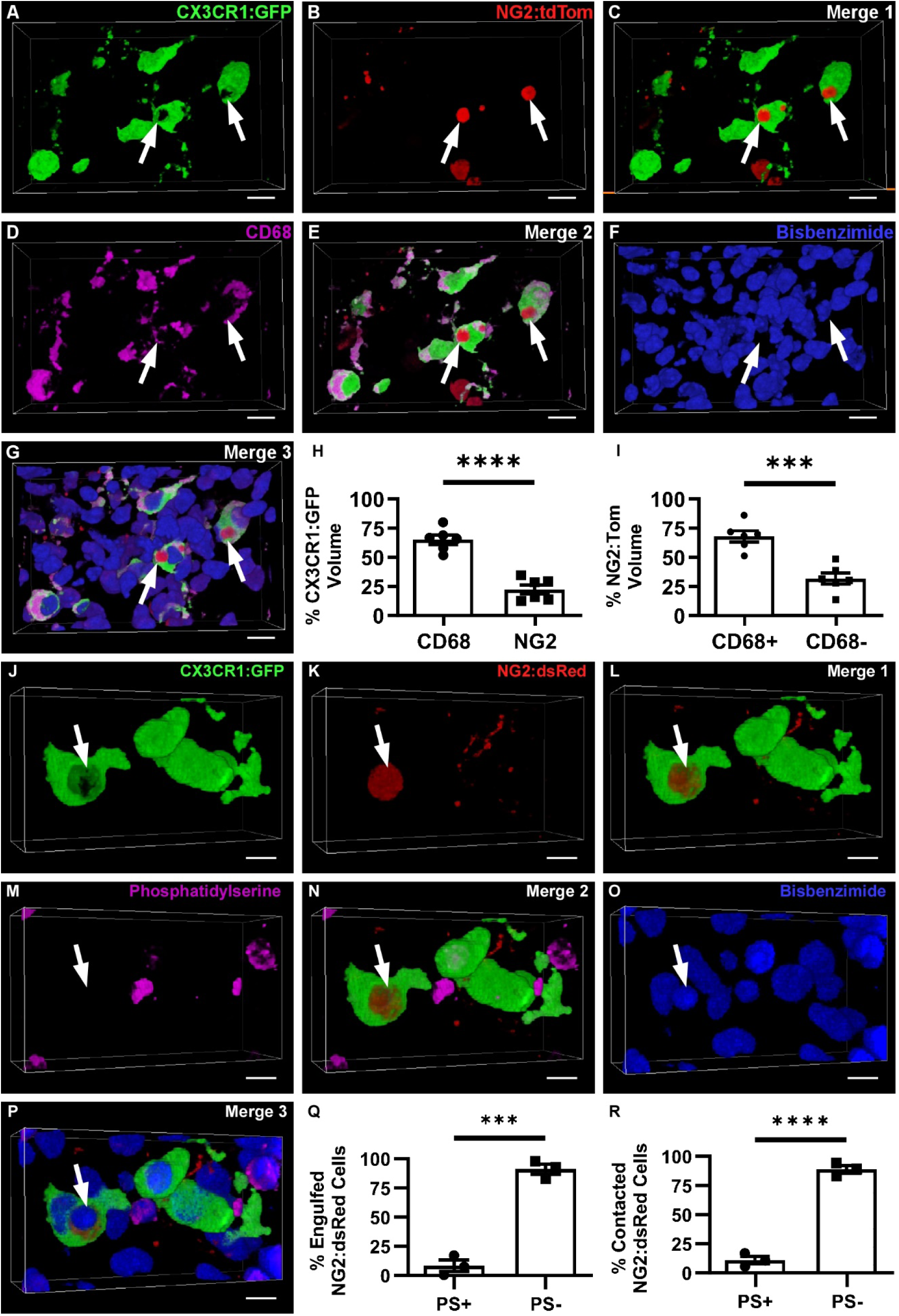
OPCs phagocytosed by microglia in the cerebellar white matter at P7 are localized to the lysosome and do not express phosphatidylserine, a marker of cell stress. 3D confocal reconstruction of CX3CR1:GFP microglia (A, green) engulfing NG2:Tom OPCs (B, red, Merge 1 in C). Staining for the CD68 lysosome (D, pink) colocalized with engulfed OPCs (E, Merge 2). Bisbenzimide was used to identify nuclear staining (F, blue) and colocalized with OPCs engulfed within the CD68 lysosome of microglia (G, Merge 3, video in supplemental data movie S2. Scale bars = 50 µm. (H) Percentage of microglial volume containing the CD68+ lysosome (mean = 65.19; SEM = 3.93) and NG2 (mean = 22.5; SEM = 3.83), \*\*\*\**P* < 0.0001. (I) Percentage of phagocytosed OPCs that colocalize (mean = 68.09; SEM = 4.72) or do not colocalize (mean = 31.91; SEM = 4.72) with CD68 staining, \*\*\**P* = 0.0003. Data from *n* = 6 animals, 4 z-stacks per animal from 2 sections. White arrow in 3D confocal z-stack indicates microglia (J) engulfing an OPC (K), shown in L (Merge 1). To evaluate cellular stress, phosphatidylserine (M) was colocalized with engulfed OPCs (N, Merge 2). Bisbenzimide nuclear staining (O) was colocalized with engulfed OPCs (P, Merge 3). Scale bars, 5 μm. (Q) Percentage of engulfed OPCs that are PS+ (mean = 8.300%, SEM = 4.807%) or PS– (mean = 91.15%, SEM = 4.36%), \*\*\**P* = 0.0002. (R) Percentage of contacted OPCs that are PS+ (mean = 11.03%, SEM = 3.252%) or PS– (mean = 88.95%, SEM = 3.276%), ****P < 0.0001. Data from *n* = 3 mice, 4 z-stacks per animal from 2 sections. Bar graphs ± SEM.

### OPCs being engulfed or contacted by microglia do not express markers of cell stress

The majority of studies exploring microglia refinement of synapses or neural progenitor cells during normal physiological conditions have shown evidence for cell death mechanisms that trigger microglia phagocytosis^37–39^. Phosphatidylserine (PS), is a phospholipid typically seen within the inner leaflet of the plasma membrane, but is exposed on the cell membrane and acts as an “eat me” signal during both apoptotic and non-apoptotic forms of cell death^40–43^. To evaluate if cell stress triggers microglia phagocytosis of OPCs in the cerebellar white matter, immunofluorescence was performed using anti-PS. To avoid any untoward effects of tamoxifen administration, NG2dsRed mice crossed with CX3CR1:GFP mice were used. Confocal 3D reconstructions of the cerebellar white matter at P7 show microglia (green, Fig. 3J) engulfing OPCs (red, Fig. 3K, merge in 3L) colocalized with anti-PS (pink, Fig. 3M, merge in N) with bisbenzimide staining (blue, Fig 3O, merge in P). Approximately, 8.30 % ± 4.80 of OPCs being engulfed expressed PS, whereas a statistically larger proportion of OPCs being engulfed did not express PS (Fig. 3Q, 91.15% ± 4.37). Similar results were observed when OPCs were contacted by microglia rather than engulfed (Fig. 3R). Of note, there were cells in the white matter that were positive for PS, but not NG2dsRed. During this developmental time period astrocytes are present as well as migrating interneurons^44^. These data suggest that the majority of OPCs being contacted or engulfed by microglia within the cerebellar white matter are viable.

### Fractalkine receptor-deficient microglia exhibit a reduction in OPC engulfment and increased number of mature oligodendrocytes

To evaluate the impact of blocking microglia engulfment of OPCs on oligodendrocyte (OL) number and maturation in the cerebellar white matter, fractalkine receptor-deficient mice (CX3CR1^mut/mut^:GFP, referred to as KO) were used to modulate microglia phagocytosis. The fractalkine-receptor is only expressed on microglia in the CNS, and fractalkine receptor-deficient mice exhibited a reduction in synaptic pruning during early postnatal development^29,45,46^. The fractalkine ligand is developmentally regulated on OPCs^47^ and we published that fractalkine receptor signaling decreased microglia engulfment of OPCs in the developing corpus callosum^25^. Representative 3D confocal reconstructions from WT (Fig. 4A) and KO (Fig. 4B) mice show a marked decrease in OPC engulfment at P7 in the cerebellar white matter. Quantification of microglia engulfment of OPCs showed that KO mice have a statistically significant reduction in engulfment compared to WT (Fig. 4C). Microglia contacting of OPCs was not statistically significant (Fig. 4D). There was no change in the number (Fig. 4E) or volume (Fig. 4F) of microglia in WT compared to KO mice. Taken together, these data suggest that fractalkine-receptor signaling on amoeboid microglia regulates engulfment of OPCs during early postnatal cerebellar white matter development.

**Figure 4.**
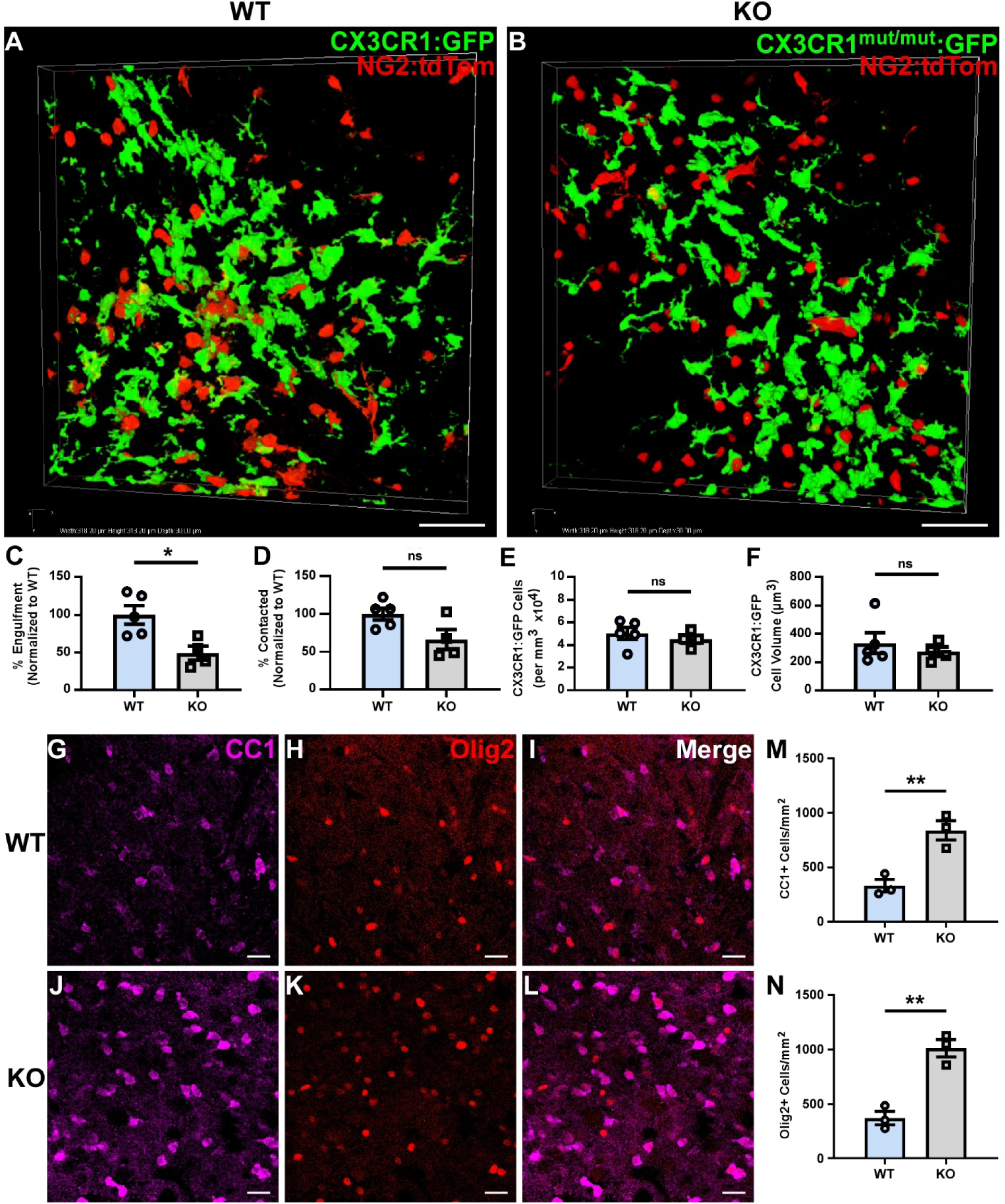
Fractalkine receptor-deficient mice show a reduction in microglia engulfment of OPCs in the cerebellar white matter at P7 and increased number of mature OLs at P30. Representative 3D reconstructed images of CX3CR1:GFP (A, WT) and CX3CR1:GFP^mut/mut^(B, KO) interactions with NG2:Tom OPCs, scale bars = 10 µm. KO mice exhibited a reduction in microglial engulfment of OPCs, characterized as 15% or more of NG2:Tom OPC volume inside microglia (C, \**P* = 0.0158). No significant changes in microglia contacting OPCs were found in WT compared to KO, characterized as less than 15% of NG2:Tom OPC volume inside microglia (D, *P* = 0.0523). There was no change in the number (E, \**P* = 0.4545) or volume (F, \**P* = 0.5095) of microglia between KO and WT mice. *n* = 5 WT and 4 KO mice, 2 sections per animal, 4 z-stacks per animal. Representative images of WT and KO mice immunostained for CC1 (G,J) and Olig2+ (H,K) colocalized in (I, L), respectively. Scale bars, 25 µm. There was a significant increase in the number of Olig2+ (M, \*\**P* = 0.003) and CC1+ (N, \*\**P* = 0.0082) OLs in KO mice. *n* = 3 WT and 3 KO mice, 3 images per animal from 3 sections. Statistical differences were determined using two-tailed unpaired *t* test. Bar graphs ± SEM.

Given that KO mice demonstrated a significant reduction in OPC engulfment and the majority of OPCs engulfed by microglia did not express PS and thus appeared viable, the potential effect on OL number at P30 was explored. Immunofluorescence staining for CC1 (mature OL marker) was performed in conjunction Olig2 (OL lineage marker) to quantify the number of mature OLs as well as the total number of OLs in cerebellar white matter at P30 (Fig. 4G-L). Cell counting revealed KO mice had significantly more CC1+ (Fig. 4M) and Olig2+ OLs (Fig. 4N) in the cerebellar white matter compared to WT mice. There was no difference in the number of NG2:Tom+ OPCs in the corpus callosum at P7 supporting the idea that OPC number was unchanged before engulfment (Fig. S1). These data are consistent with our results indicating that OPCs are viable before engulfment, and that blocking engulfment of OPCs resulted in their survival (i.e. increased numbers) and subsequent maturation. Although death of cells or synapses before removal by microglia is a generally accepted mechanism, the idea that blocking microglia phagocytosis prevents death of the target cell is termed primary phagocytosis or phagoptosis Several studies have documented that during development neural cells are viable before engulfment by microglia, and blocking engulfment resulted in their increased numbers^48,49^. consistent with findings in our study.

### Fractalkine receptor-deficient mice show reduced myelin thickness with no change in axon number

To further explore how changes in OL number and maturation affect myelin thickness, electron microscopy was performed in the cerebellar white matter of P30 KO and WT mice. Images of WT (Fig. 5A-B) and KO (Fig. 5C-D) mice were analyzed for changes in myelin thickness using MyelTracer 2.0, a semi-automated program for evaluating myelin and axonal diameter that was found to be consistent with previously published methods using ImageJ^25^ (Fig. S2-S3). KO mice had a decrease in myelin thickness compared to WT (Fig. 5A-D), which was represented by an increase in g ratio (axon diameter divided by axon plus myelin diameter, Fig. 5E). Individual g ratios were plotted as a function of their corresponding axon diameter, which revealed a statistically significant difference between KO and WT mice (Fig. 5F). Moreover, there was a statistically significant shift in the distribution of g ratios towards larger values in KO compared to WT mice (Fig. 5G). There was no change in axon number across both groups (Fig. 5H), however there was a statistically significant increase in the percentage of unmyelinated axons (Fig. 5I) and a corresponding decrease in the percentage of myelinated axons (Fig. 5J) in KO mice. These data suggest that microglia engulf OPCs during development as part of a homeostatic mechanism to regulate the proper number of OLs needed to promote efficient myelination of axons within the cerebellar white matter.

**Figure 5.**
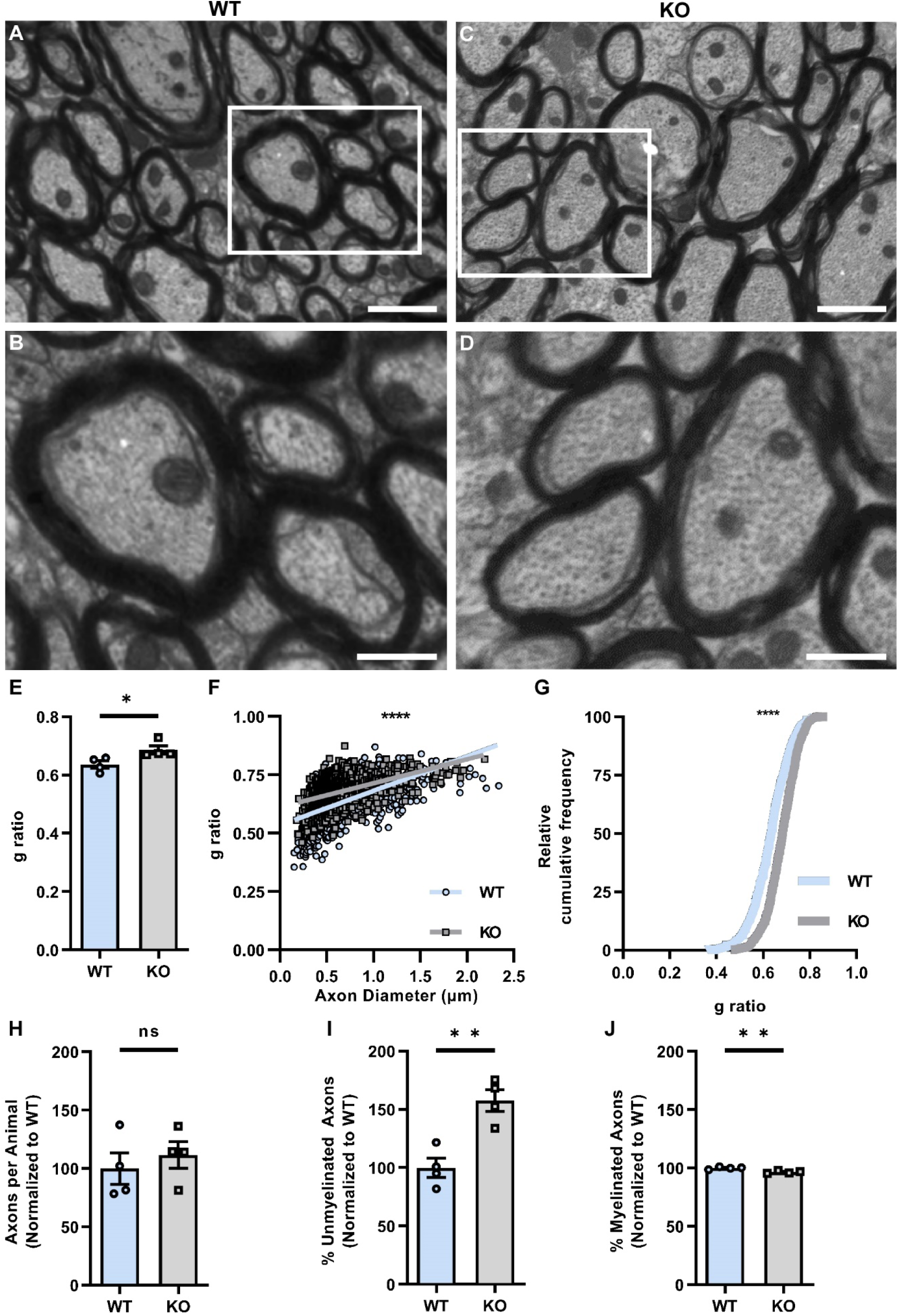
Impaired engulfment of OPCs in fractalkine receptor-deficient mice leads to impaired myelin thickness at P30 in the cerebellar white matter. Representative electron microscopy images of CX3CR1:GFP (WT, A) and CX3CR1^mut/mut^:GFP (KO, C) cerebellar white matter at P30. Scalebars = 2 µm. White boxes represent magnified areas in B and D, respectively. Scalebars, 1 µm. (E) There was an increase in g-ratio (axon diameter divided by axon plus myelin diameter) reflecting a decrease in myelin thickness in KO compared to WT mice (\**P* = 0.0357). (F) Scatter plot of g ratio for each axon diameter was statistically significant different between WT and KO. (\*\*\*\**P* = 0.0001, analysis of covariance). Relative cumulative frequency of g ratios (G) was significantly different between WT and KO mice (\*\*\*\**P* = 0.0001, Kolmogorov-Smirnov test). There was no significant difference in the number of axons (H, \**P* = 0.5343), however KO mice showed a significant increase in the percentage of unmyelinated axons (I, \*\**P* = 0.0035) and a subsequent decrease in the percentage of myelinated axons (J, \*\**P* = 0.0044). Statistical differences were determined using a two-tailed *t* test unless where indicated, *n* = 4 mice per genotype, approximately 1,000 axons per genotype, from 2-8 images per mouse. Bar graphs ± SEM.

## Discussion

Proper myelin formation is not only associated with ensuring proper conduction of action potentials, but also providing metabolic support to axons^50,51^. This suggests that impairments in cerebellar white matter myelination could have grave consequences for the integrity and function of Purkinje neurons whose axons are housed in the cerebellar white matter. Purkinje neurons not only integrate all incoming information to the cerebellum, but are also the only output neurons for the cerebellum including sending information to the cerebral cortex. Here, we show that fractalkine receptor-deficiency in microglia decreases microglia engulfment of OPCs that ultimately results in reduced myelin thickness in the cerebellar white matter at P30. These findings have important implications for neurodevelopmental disorders such as autism spectrum disorder that are associated with variants in the fractalkine receptor that alter its function^34^. Several clinical imaging studies have documented cerebellar white matter reduction in patients with autism spectrum disorder^30–33^. Fractalkine receptor KO mice also exhibit deficits in developmental synaptic pruning by microglia^29^. The concept of deficits in synaptic pruning leading to overproduction of synapses and impaired communication has been confirmed in postmortem autism brains where increased density of synapses has been reported^52^. Fractalkine receptor-deficient mice exhibit impairments in social interaction, increased repetitive behaviors, and motor dsyfunction^45,53^ that are considered hallmarks of autism spectrum disorder and other neurodevelopmental disorders. Taken together these data suggest that deficits in microglia-mediated pruning in autism may not be limited to synaptic pruning, but also pruning of OPCs, which occurs during similar developmental time periods.

Here, we show that refinement of cerebellar white matter occurs during early postnatal development concurrent with the corpus callosum as we previously published^36^. This synchronization of amoeboid microglial phagocytosis of OPCs in both cortical and cerebellar white matter and the resulting hypomyelination recapitulated in these regions in fractalkine receptor KO mice posits an unexplored link for white matter alterations in neurodevelopmental disorders. Furthermore, the known impact of proper myelination on neuronal health^50,51^ could also contribute to cortical and cerebellar neuronal dysfunction. The predominant population of axons in the corpus callosum are cortical layer II/III, and V pyramidal neurons that play significant roles in higher order cognitive processing^54^. Alterations in pyramidal as well as Purkinje neurons are reported in developmental disabilities^55,56^. Phagocytosis of OPCs by microglia is restricted to the white matter tracks in both the cortex and cerebellum that house the axons of the aforementioned neurons, supporting an important homeostatic process necessary for brain development. Disruptions in both cerebral and cerebellar white matter tracks in autism spectrum disorder^30–33^ and an association with genetic variants in the fractalkine receptor^34^, suggest that altered microglia refinement of white matter during early postnatal development may be an underexplored mechanism for neurodevelopmental disorders.

The presence of amoeboid microglia has been described in human perinatal white matter^57,58^, a time period that correlates with the maturational stage of rodent white matter studied here^59^. While neurodevelopmental and neuropsychiatric disorders are associated with genetic risk factors for microglia and alterations in myelin^16,17^, leukodystrophies also share this common link. Leukodystrophies are hypomyelinating developmental disorders that have mutations in genes important for microglial proliferation and phagocytosis such as the *Csf1r* gene^60–64^. Similarly, Nasu-Hakola disease, characterized by early-onset dementia with sclerosing leukoencephalopathy and bone cysts, is caused by mutations of the triggering receptor expressed on myeloid cells or its adapter protein, tyrosine kinase-binding protein^65^, a protein complex only expressed on microglia^66^. These clinical findings in conjunction with the preliminary data presented here highlight the role of microglia in white matter formation, which may have long-term implications for axonal health affecting neuronal function.

While proliferation of OPCs appears to be a part of normal white matter development that require refinement by microglia, it would be of interest to understand if this process is lacking in adult demyelinating diseases where OPCs also proliferate but remyelination is not robust. Interestingly, several studies have documented the important role of microglia phagocytosis in the removal of myelin debris^67–69^, as a necessary mechanism for remyelination including the fractalkine receptor ^70^ (review^71^). It appears that microglia in both demyelinating and neurodegenerative disease states exhibit a disease-associated phenotype^72^ that may alter their normal homeostatic processes including refinement of the OPC pool.

We have shown that amoeboid microglia are only found in the white matter during a restricted window of development. These microglia do have a unique phagocytic transcriptional profile^73,74^, which we have previously reported phagocytose OPCs in the corpus callosum^36^ and, here, in the cerebellum. Overall, these findings suggest that the infiltration of amoeboid microglia is restricted to white matter regions, and serves a homeostatic function to refine OPCs that is conserved in a spatiotemporal manner to modulate myelin formation.

## Acknowledgments

We would like to thank Judith Drazba, Director of the Cleveland Clinic Imaging core, and Mei Yin, resource specialist, for assistance with electron microscopy.

## Funding

This study was supported by National Multiple Sclerosis Society RG 4587-A-1, National Science Foundation 1648822, National Eye Institute RO1EY025687, and the Mike L. Jezdimir Transverse Myelitis Foundation.

## Author contributions

T.M.D, A.D.N, and J.S. conceived the study and designed the experiments. M.K.C. and A.D.N. completed the engulfment analysis. M.K.C and J.S. performed the EM analyses. M.K.C., J.S., M.I., and G.M.M. contributed to immunocytochemistry. E.S. and J.S. conceived the idea and wrote the code for MyelTracer 2.0. T.M.D., M.K.C. and J.S. drafted and edited the manuscript.

## Competing interests

The authors declare that they have no competing interests.

## Data and materials availability

All data needed to evaluate the conclusions in the paper are present in the paper and/or the Supplementary Materials. Additional data related to this paper may be requested from the authors.

**Fig. S1.**
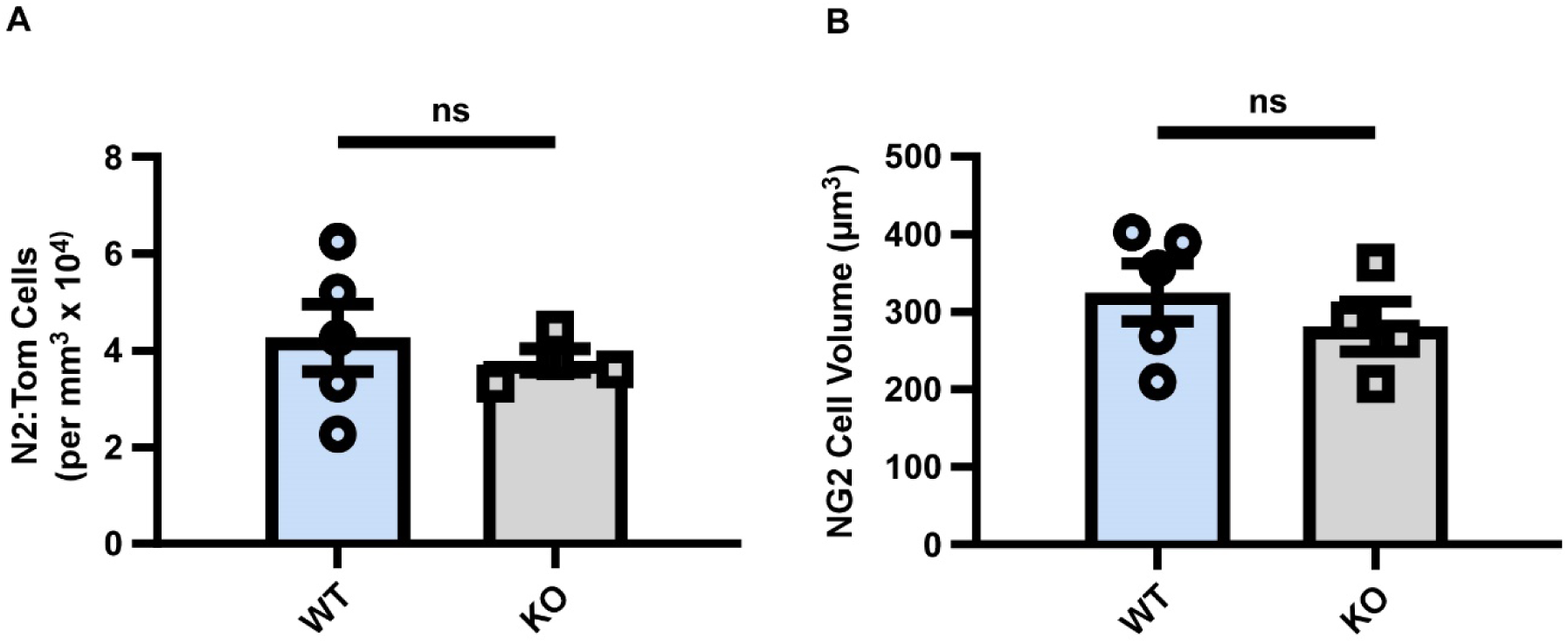
Fractalkine receptor-deficient mice do not exhibit a change in the number of OPCs in cerebellar white matter at P7. NG2:Tom OPCs were counted in CX3CR1:GFP (WT) and CX3CR1:GFP^mut/mut^ (KO) mice, representative images in Fig. 4A-B. There was no change in the number (A, \**P* = 0.5825) or volume (B, \**P* = 0.4129) of NG2+ OPCs between KO and WT mice in P7 cerebellum, *n* = 5 WT and 4 KO, 4 z-stacks per animal from 2 sections. Bar graphs ± SEM.

**Figure S2.**
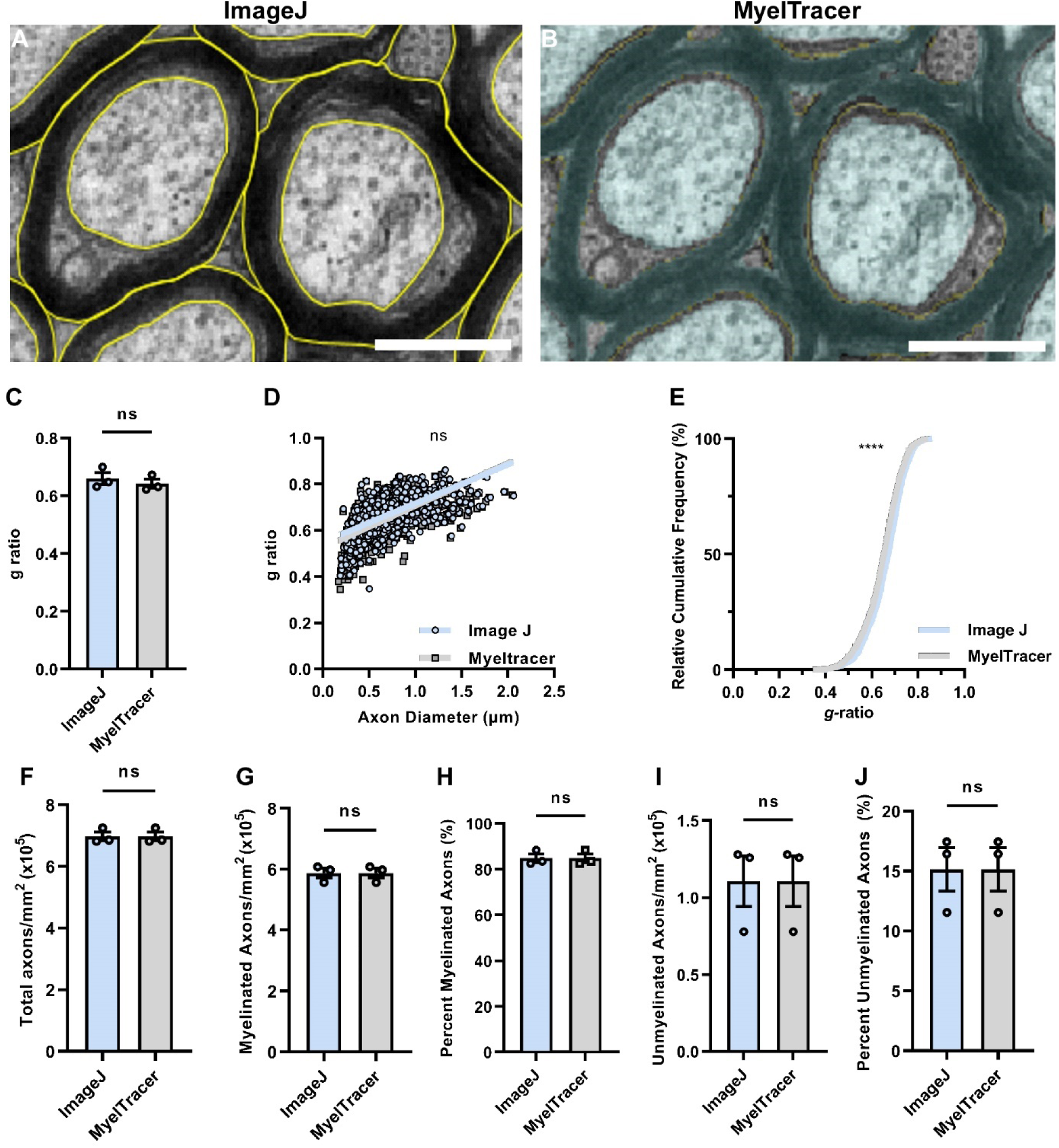
Comparative analysis between manual tracing using ImageJ and semi-automated segmentation using MyelTracer 2.0. Representative electron microscopy images of manual (A, Image J) versus semi-automated (B, MyelTracer 2.0) tracing of axons in the optic nerve. Scale bars = 0.5 µm. There was no change in g ratios using ImageJ compared to MyelTracer 2.0 (C, *P* = 0.5304, student’s unpaired t-test). Scatter plot of g ratio for each axon diameter showed no statistical difference (D, *P* = 0.0952, analysis of covariance). Relative cumulative frequency distribution of g ratios did show slightly larger g ratio measurements in MyelTracer 2.0 analyzed images (E, *P* <0.0001, Kolmogorov-Smirnov test), explanation in Fig. S3. There was no change in total number of axons (F, *P* > 0.9999), myelinated axon number (G, *P* > 0.9999), percent myelinated axons (H, *P* > 0.9999), unmyelinated axon number (I, *P* > 0.9999), and percent myelinated axons (J, *P* > 0.9999), *n* = 3 mice per group, 6 images per mouse. Bar graphs ± SEM.

**Figure S3.**
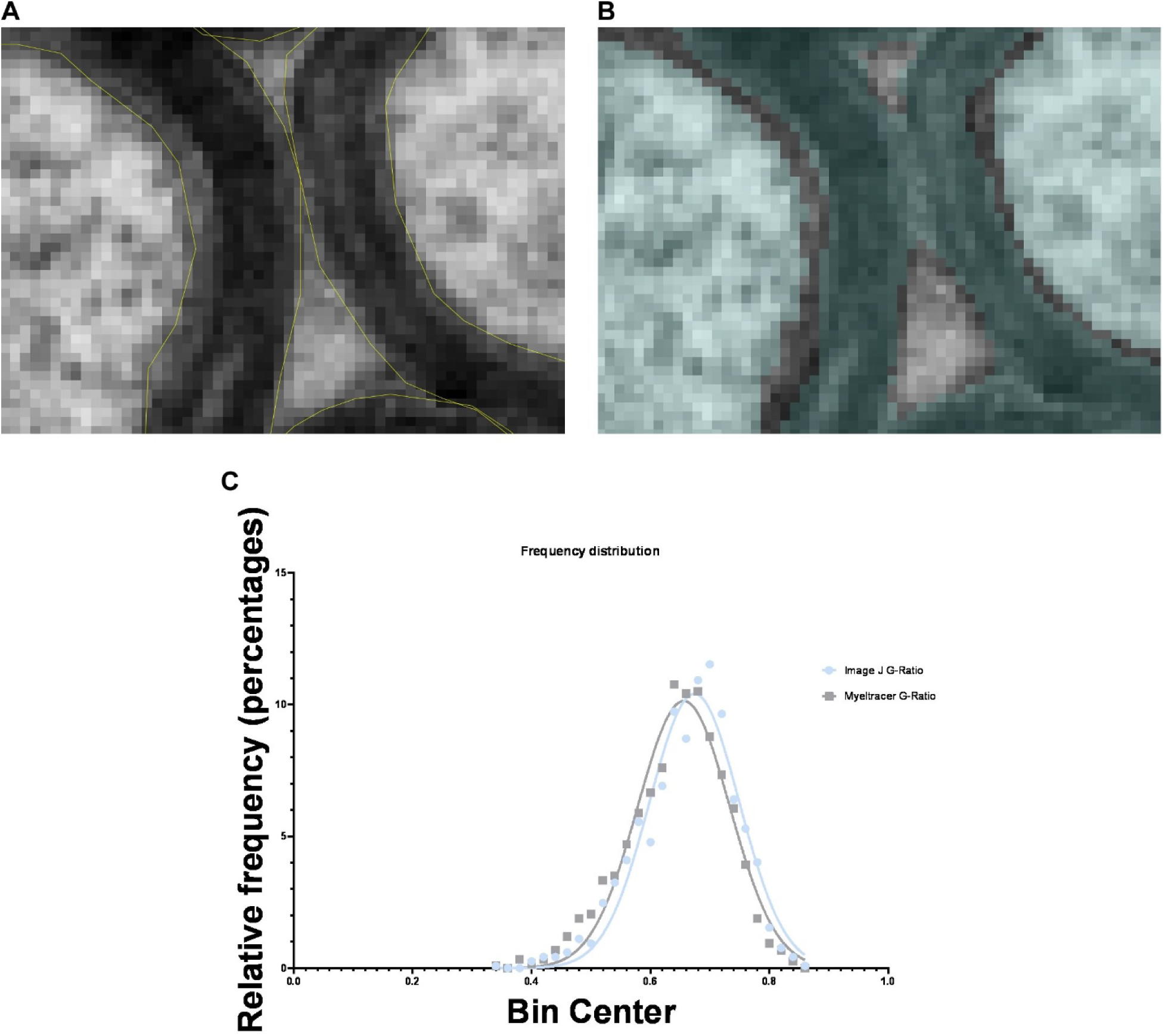
Representative images of border tracing using ImageJ (A) and MyelTracer 2.0 (B). Images were digitally zoomed to magnify the borders of axon and myelin sheath. Non-linear regression of g ratio frequency distributions shows a reduction in variability of g ratio measured using MyelTracer 2.0 software (C), *R*^2^: IJ = 0.9508, MT = 0.9663, Sum of Squares: IJ = 18.30, MT = 11.61.

## Supplementary Movies

**Movie S1 Amoeboid microglia encapsulate OPCs in the cerebellar white matter at P7.**

**Movie S2 Phagocytosed OPCs are identified within lysosomes of amoeboid microglia**

## Methods

### RESOURCE AVAILABILITY

#### Lead Contact

Further information and requests for resources and reagents should be directed to and will be fulfilled by the Lead Contact, Tara DeSilva.

#### Materials Availability

*This study did not generate new unique reagents*.

#### Data and Code Availability

- All data reported in this paper will be shared by the lead contact upon request.
- The original code/software is freely available online at https://github.com/HarrisonAllen/ MyelTracer. The updated code (MyelTracer 2.0) used throughout the paper is freely available online at https://github.com/jackslate45/MyelTracer. The code is available as Extended Data 1.
- Any additional information required to reanalyze the data reported in this paper is available from the lead contact upon request.

### EXPERIMENTAL MODEL AND SUBJECT DETAILS

#### Animals

NG2:Tom (NG2-creER:tdTomato Jax 008538 backcrossed with Jax 007914), NG2dsRed (Jax 008241), and CX3CR1:GFP (Jax 005582) were received from the Jackson Laboratory (Bar Harbor, ME). Homozygous CX3CR1^mut/mut^ mice are referred to as KO throughout the text, and CX3CR1:GFP mice, which have heterozygous expression of CX3CR1, are referred to as WT. Both male and female mice were used for all experiments as well as littermate controls. Animals were housed and treated in accordance with the United States National Institutes of Health guidelines for the humane treatment of animals and experiments were approved by the Institutional Animal Care and Use Committee of Cleveland Clinic. Mice were housed in individually ventilated cages under a standard 12 hour light/dark cycle with 18% protein standard chow and sterilized water provided *ad libitum*. Each cage contained 2-5 mice. Cages contained pulped cotton fiber nestlets for enrichment. Mice were weaned at 3 weeks of age.

### METHOD DETAILS

#### Tamoxifen induction

NG2:Tom mice received intraperitoneal injections (i.p.) with 10 mg/mL tamoxifen (Sigma-Aldrich, St. Louis, MO; T5648) in 30 μL peanut oil. For pups taken at P4, tamoxifen i.p. injections were administered at P1. For pups taken at P7 and older, tamoxifen was injected at P3.

#### Perfusion and preparation of mice brains for immunofluorescence

Transcardiac perfusion was administered with phosphate buffered saline (PBS) followed by 4% paraformaldehyde (PFA). Brains were collected and post-fixed for 24 hours in 4% PFA at 4°C, then moved to 30% sucrose for 48 hours at 4°C. Brains were then embedded in 1 part 30% sucrose in PBS and 2 parts optimal cutting temperature compound (OCT) (Fisher HealthCare, Waltham, MA) and stored at –80°C. Embedded brains were sectioned into 15 μm thick sagittal slices with a Leica cryostat (Leica Biosystems, Nussloch, Germany).

#### Immunofluorescent staining

Embedded brains were stained as recommended by the antibody manufacturer. Sections were washed with PBS and blocked with a 5% serum corresponding to the host of the secondary antibody with 0.3% Triton X-100 for 1 hour at room temperature. Antigens expressed on the surface of the cell membrane (i.e. phosphatidylserine) were stained without Triton X-100. Primary antibodies were diluted in blocking buffer and incubated overnight at 4°C and subsequently washed before incubation with secondary antibodies for 1 hour at room temperature. Following a wash step, the sections were coverslipped with Fluoromount-G (Southern Biotech, Birmingham, AL; 0100-01) containing bisbenzimide (1:1,000; Invitrogen, Carlsbad, CA; H3569). Primary antibodies used in this study included rat anti-mouse CD68 (1:200; Biorad, mca1957ga), mouse anti-phosphatidylserine (1:200 without Triton, Millipore, 05719), mouse anti-CC1 (1:100 free-floating; Millipore, OP80), and rabbit anti-Olig2 (1:200 free-floating; Millipore, ab9610). Secondary antibodies included Alexa Fluor 647 goat anti-mouse IgG (1:1,000; Invitrogen, A21235) and Alexa Fluor 594 goat anti-rabbit IgG (1:500; Invitrogen, A11012).

#### Imaging

Images were acquired using a Nikon C2 confocal microscope with NIS-Elements software (Nikon, Melville, NY). Z-stacks for 3D reconstructions were captured at 15 μm thickness using a 0.3 μm step size with a Plan Fluor 40X/1.30 NA (oil immersion) objective. Images were captured from the medial portion of the cerebellar white matter adjacent to the fourth ventricle referred to as the trunk of arbor vitae.

#### Electron Microscopy

At P30, mice were injected with 100 µL of 1% lidocaine in the left lateral ventricle prior to transcardiac perfusion with 4% PFA and 2% glutaraldehyde in 0.1M cacodylate buffer at a flow rate of 10 mL/min for 10 minutes. Brains were collected and sectioned into 500 µm-thick sagittal sections with a vibrating microtome (Leica, Heidelberg, Germany). The most medial portion of the cerebellar white matter tracts was dissected from the cerebellum and processed by the Cleveland Clinic Electron Microscopy Core. The tissue blocks were post-fixed with 1% osmium tetroxide in water, stained with 1% uranyl acetate in maleate buffer, then dehydrated with a mixture of ethanol and propylene oxide. Following the dehydration step, tissue was embedded in Pure Eponate 12 resin (Ted Pella Inc., Redding, CA). The tissue was cut into ultra-thin sections, stained with uranyl acetate and lead citrate, and imaged with a transmission electron microscope (FEI Company, Hillsboro, OR).

### Quantification and Statistical Analysis

#### Engulfment analysis

Methods from our published study were used to analyze microglial engulfment of OPCs ^25,36^. Confocal 3D z-stacks acquired on Nikon were imported into Imaris software (Bitplane, Zurich, Switzerland). Surface rendering for microglia and OPCs was conducted using 0.1 μm and 0.4 μm smoothing, respectively. Absolute thresholding was performed by a researcher blinded to group identity and surfaces smaller than 10 μm^3^ were filtered out. Surface renders were separated or unified using bisbenzimide as the marker of the cell nucleus. Pericytes were excluded by filtering out any surfaces with a volume less than 10 μm^3^ and sphericity less than 0.49. These criteria were determined based on the size of pericytes on blood vessels (Peppiatt et al., 2006). Following the completion of surface renders, the surface-surface coloc XTension software available through Imaris was used to create a new surface channel that identified overlapping voxels between microglial and OPC surfaces. Microglia to OPC surface volumes were divided by their corresponding OPC volumes to determine the percentage of OPC internalized by the microglia. Engulfment was categorized as ≥15% of NG2:Tom OPC volume internalized by microglia and contacting as ≤15% of NG2:Tom OPC volume internalized by microglia.

#### Quantification of CD68 Colocalization

Sections from P7 CX3CR1:GFP;NG2:Tom mice were stained for CD68 and analyzed within the cerebellar white matter. A total of 4 confocal z-stacks were taken per animal from 2 sections approximately 150 µm apart. The z-stacks were taken at 40X magnification with 200 μm x 200 μm x 15 μm fields. Surface renders were created for each channel through manual thresholding using the Imaris software (Bitplane, Zurich, Switzerland). The percentage of microglial volume that contained a CD68+ lysosomal staining and NG2:Tom+ cell volume was quantified as described in studies on synapse engulfment (Schafer et al., 2012). NG2:Tom+ cells engulfed by microglia were categorized as being positive or negative for CD68 staining.

#### Quantification of cellular stress

Sections from P7 CX3CR1:GFP; NG2:dsRed mice were stained with antibodies generated against phosphatidylserine and analyzed within the cerebellar white matter. A total of 4 z stacks per animal were collected at 40x magnification with 200 μm x 200 μm x 15 μm fields from 2 sections approximately 150 µm apart. Imaris software (Bitplane, Zurich, Switzerland) was used to analyze OPC contact, or engulfment as described in the “Engulfment analysis” method. Engulfed or contacted cells were further analyzed as being immunopositive or negative for phosphatidylserine staining.

#### Quantification of engulfment in CX3CR1 KO mice

Sections from P7 CX3CR1:GFP;NG2:Tom (WT) and CX3CR1^mut/mut^:GFP;NG2:Tom (KO) mice were collected and imaged with 4 z-stacks per animal from 2 sections approximately 150 µm apart. Confocal z-stacks were taken at 40X magnification with 0.3 μm step and 200 μm x 200 μm x 15 μm fields. Images were analyzed for OPC engulfment or contact by microglia using Imaris software (Bitplane, Zurich, Switzerland) as described in the “Engulfment analysis” method.

#### Quantification of oligodendrocyte number at P30

Cell counts were assessed by a researcher blinded to genotype on sections from P30 CX3CR1:GFP;NG2:Tom (WT) and CX3CR1^mut/mut^:GFP;NG2:Tom (KO) mice stained for CC1 and Olig2. CC1 is a marker of mature oligodendrocytes, whereas Olig2 is a marker of all oligodendrocytes. Sagittal sections of the cerebellar white matter were analyzed with 3 images per animal from 3 sections approximately 150 µm apart. Images were taken at 20X magnification with 200 μm x 200 μm fields. Cells were counted as positive for CC1 or Olig2 using the cell counter plugin with free NIH software (ImageJ). All cells counted were confirmed to have a bisbenzimide-positive nucleus.

### MyelTracer update to MyelTracer 2.0 (https://github.com/jackslate45/MyelTracer)

MyelTracer, being an open-source platform, was upgraded to MyelTracer 2.0, by adding features to assist in the analysis of electron microscopy images. First, three additional counters were added for tracking cellular structures independent from the original unmyelinated and myelinated trackers. Next, three additional miscellaneous selection tools were added, each distinguished by a different color, to measure and identify cellular structures distinct from the original selection tool. Measuring axon diameter divided by outer myelin diameter has been the standard in the field for measuring g ratios. This calculation was added to MyelTracer 2, in addition to, the inner myelin divided by outer myelin selection tool developed as part of original MyelTracer platform. Also distinct IDs were made for the additional selection and counting tools created in the excel file so that each measurement can be accurately associated with the proper tool ID.

#### Analysis of Electron Microscopy Images

To evaluate changes in myelination, tissue from the cerebellar white matter tracts of P30 CX3CR1:GFP (WT) and CX3CR1^mut/mut^:GFP (KO) mice was imaged using electron microscopy. Two to eight images were captured per mouse at either 6800X or 9300X magnification and were used to measure g ratios and axon numbers by a researcher blinded to genotype. Using MyelTracer 2.0, axons were counted as myelinated or unmyelinated. Only axons containing microtubules and neurofilaments were included. Myelinated axons were identified based on the presence of dark bands surrounding axons. MyelTracer 2.0 was used to semi-automatically quantify g ratios in a threshold-dependent manner. The MyelTracer software used to evaluate the electron microscopy images was validated against the manual tracing method commonly used in ImageJ. Manual tracing in ImageJ was performed using the polygon tool to trace axons first, then the outer border of the myelin sheaths (Fig. S2A). This was compared to the semi-automated analysis of MyelTracer (Fig. S2B) using 6 images from 3 mice at postnatal day 90. MyelTracer has previously been shown to analyze EM data for g-ratio analysis with similar output to manual tracing with a ∼60% reduction in time spent on analysis^75^. Similarly, our data showed no change in g ratio (Fig. S2C). Using a scatter plot to evaluate each g ratio with its corresponding axon diameter there was also no significant difference between ImageJ and MyelTracer (Fig. S2D). There was a statistically significant difference in the relative cumulative frequency across g ratios (Fig. S2E). However, there was no statisital difference in the total axons (Fig. S2F), myelinated axons (Fig. S2G), percent myelinated axons (Fig. S2H), unmyelinated axons (Fig. S2I), or percent unmyelinated axons (Fig. S2J) between ImageJ and MyelTracer. These data indicated a slightly larger measured myelin sheath using MyelTracer compared to Image J (Fig. S2E). Regression analysis on this distribution data showed that MyelTracer reduced variability in the measures of g ratio as compared to manual tracing (*R*^2^: IJ = 0.9512, MT = 0.9660, Sum of Squares: IJ = 18.16, MT = 11.74. (Fig. S3). The ability of MyelTracer to draw single pixel-based borders and the use of thresholding allowed for a more accurate-unbiased measurement of the border of the myelin sheath by relying more on the grayscale value of the pixel and not the discernment of the measurer. The small decrease in the g ratio measure more accurately represents the border of the myelin and is more accurately traced to the contours of the pixel than is possible using manual tracing methods.

### Statistical analyses

All statistical analyses were performed using GraphPad Prism software version 9.0 (GraphPad Software, La Jolla, CA) except the ANCOVA used for EM analysis was performed in R. Analyses including p values are reported where indicated in the figure legends and results.

## Notes

https://github.com/jackslate45/MyelTracer

### Competing Interest Statement

TMD serves on the board of directors for Coeptis Therapeutics. All other authors declare no competing interests.

## REFERENCES

1. Itō, M. The cerebellum and neural control, (Raven press, 1984).

2. Carta, I., Chen, C.H., Schott, A.L., Dorizan, S. & Khodakhah, K. Cerebellar modulation of the reward circuitry and social behavior. Science 363, eaav0581 (2019).

3. Carleton, S.C. & Carpenter, M.B. Distribution of primary vestibular fibers in the brainstem and cerebellum of the monkey. Brain research 294, 281–298 (1984).

4. Simpson, J., Precht, W. & Llinás, R. Sensory separation in climbing and mossy fiber inputs to cat vestibulocerebellum. Pflügers Archiv 351, 183–193 (1974).

5. Ha, H. & Liu, C.N. Cell origin of the ventral spinocerebellar tract. Journal of Comparative Neurology 133, 185–205 (1968).

6. Matsushita, M. & Hosoya, Y. Cells of origin of the spinocerebellar tract in the rat, studied with the method of retrograde transport of horseradish peroxidase. Brain research 173, 185–200 (1979).

7. Lundberg, A. & Weight, F. Functional organization of connexions to the ventral spinocerebellar tract. Experimental brain research 12, 295–316 (1971).

8. Kamali, A., Kramer, L.A., Frye, R.E., Butler, I.J. & Hasan, K.M. Diffusion tensor tractography of the human brain cortico-ponto-cerebellar pathways: a quantitative preliminary study. Journal of Magnetic Resonance Imaging 32, 809–817 (2010).

9. Palesi, F., et al. Contralateral cortico-ponto-cerebellar pathways reconstruction in humans in vivo: implications for reciprocal cerebro-cerebellar structural connectivity in motor and non-motor areas. Scientific reports 7, 12841 (2017).

10. Henschke, J.U. & Pakan, J.M. Disynaptic cerebrocerebellar pathways originating from multiple functionally distinct cortical areas. elife 9, e59148 (2020).

11. Ramnani, N., et al. The Evolution of Prefrontal Inputs to the Cortico-pontine System: Diffusion Imaging Evidence from Macaque Monkeys and Humans. Cerebral Cortex 16, 811–818 (2005).

12. Akintunde, A. & Buxton, D.F. Origins and collateralization of corticospinal, corticopontine, corticorubral and corticostriatal tracts: a multiple retrograde fluorescent tracing study. Brain research 586, 208–218 (1992).

13. Shinoda, Y., Sugiuchi, Y., Futami, T. & Izawa, R. Axon collaterals of mossy fibers from the pontine nucleus in the cerebellar dentate nucleus. Journal of neurophysiology 67, 547–560 (1992).

14. Allen, G., Azzena, G. & Ohno, T. Pontine and non-pontine pathways mediating early mossy fiber responses from sensorimotor cortex to cerebellum in the cat. Experimental Brain Research 36, 359–374 (1979).

15. De Camilli, P., Miller, P.E., Levitt, P., Walter, U. & Greengard, P. Anatomy of cerebellar Purkinje cells in the rat determined by a specific immunohistochemical marker. Neuroscience 11, 761–IN762 (1984).

16. Funfschilling, U., et al. Glycolytic oligodendrocytes maintain myelin and long-term axonal integrity. Nature 485, 517–521 (2012).

17. Lee, Y., et al. Oligodendroglia metabolically support axons and contribute to neurodegeneration. Nature 487, 443–448 (2012).

18. Gianola, S., Savio, T., Schwab, M.E. & Rossi, F. Cell-Autonomous Mechanisms and Myelin-Associated Factors Contribute to the Development of Purkinje Axon Intracortical Plexus in the Rat Cerebellum. The Journal of Neuroscience 23, 4613–4624 (2003).

19. Zagrebelsky, M., et al. Retrograde regulation of growth-associated gene expression in adult rat Purkinje cells by myelin-associated neurite growth inhibitory proteins. Journal of Neuroscience 18, 7912–7929 (1998).

20. Barron, T., Saifetiarova, J., Bhat, M.A. & Kim, J.H. Myelination of Purkinje axons is critical for resilient synaptic transmission in the deep cerebellar nucleus. Scientific Reports 8, 1022 (2018).

21. Stevens, B., et al. The classical complement cascade mediates CNS synapse elimination. Cell 131, 1164–1178 (2007).

22. Paolicelli, R.C., et al. Synaptic Pruning by Microglia Is Necessary for Normal Brain Development. Science 333, 1456–1458 (2011).

23. Hoshiko, M., Arnoux, I., Avignone, E., Yamamoto, N. & Audinat, E. Deficiency of the microglial receptor CX3CR1 impairs postnatal functional development of thalamocortical synapses in the barrel cortex. Journal of Neuroscience 32, 15106–15111 (2012).

24. Ginhoux, F., et al. Fate mapping analysis reveals that adult microglia derive from primitive macrophages. Science 330, 841–845 (2010).

25. Nemes-Baran, A.D., White, D.R. & DeSilva, T.M. Fractalkine-Dependent Microglial Pruning of Viable Oligodendrocyte Progenitor Cells Regulates Myelination. Cell reports 32, 108047 (2020).

26. Levison, S.W. & Goldman, J.E. Both oligodendrocytes and astrocytes develop from progenitors in the subventricular zone of postnatal rat forebrain. Neuron 10, 201–212 (1993).

27. Kessaris, N., et al. Competing waves of oligodendrocytes in the forebrain and postnatal elimination of an embryonic lineage. Nature neuroscience 9, 173–179 (2006).

28. Delassalle, A., et al. Regional distribution of myelin basic protein in the central nervous system of quaking, jimpy, and normal mice during development and aging. Journal of neuroscience research 6, 303–313 (1981).

29. Paolicelli, R.C., et al. Synaptic pruning by microglia is necessary for normal brain development. Science 333, 1456–1458 (2011).

30. Cheung, C., et al. White matter fractional anisotrophy differences and correlates of diagnostic symptoms in autism. J Child Psychol Psychiatry 50, 1102–1112 (2009).

31. Kleinhans, N.M., et al. Age-related abnormalities in white matter microstructure in autism spectrum disorders. Brain Res 1479, 1–16 (2012).

32. Ecker, C., et al. Brain anatomy and its relationship to behavior in adults with autism spectrum disorder: a multicenter magnetic resonance imaging study. Arch Gen Psychiatry 69, 195–209 (2012).

33. Hampson, D.R. & Blatt, G.J. Autism spectrum disorders and neuropathology of the cerebellum. Frontiers in neuroscience 9, 420 (2015).

34. Ishizuka, K., et al. Rare genetic variants in CX3CR1 and their contribution to the increased risk of schizophrenia and autism spectrum disorders. Transl Psychiatry 7, e1184 (2017).

35. Levison, S.W. & Goldman, J.E. Both oligodendrocytes and astrocytes develop from progenitors in the subventricular zone of postnatal rat forebrain. Neuron 10, 201–212 (1993).

36. Nemes-Baran, A.D. & DeSilva, T.M. Quantification of microglial contact and engulfment of oligodendrocyte progenitor cells in the rodent brain. STAR Protoc 2, 100403 (2021).

37. Sierra, A., et al. Microglia shape adult hippocampal neurogenesis through apoptosis-coupled phagocytosis. Cell stem cell 7, 483–495 (2010).

38. Schafer, D.P., et al. Microglia sculpt postnatal neural circuits in an activity and complement-dependent manner. Neuron 74, 691–705 (2012).

39. Gunner, G., et al. Sensory lesioning induces microglial synapse elimination via ADAM10 and fractalkine signaling. Nat Neurosci 22, 1075–1088 (2019).

40. Shlomovitz, I., Speir, M. & Gerlic, M. Flipping the dogma – phosphatidylserine in non-apoptotic cell death. Cell Commun Signal 17, 139 (2019).

41. de Vasconcelos, N.M., Van Opdenbosch, N., Van Gorp, H., Parthoens, E. & Lamkanfi, M. Single-cell analysis of pyroptosis dynamics reveals conserved GSDMD-mediated subcellular events that precede plasma membrane rupture. Cell Death Differ 26, 146–161 (2019).

42. Andrabi, S.A., Dawson, T.M. & Dawson, V.L. Mitochondrial and nuclear cross talk in cell death: parthanatos. Ann N Y Acad Sci 1147, 233–241 (2008).

43. Lee, J.Y., Kim, W.K., Bae, K.H., Lee, S.C. & Lee, E.W. Lipid Metabolism and Ferroptosis. Biology (Basel) 10(2021).

44. Groteklaes, A., Bonisch, C., Eiberger, B., Christ, A. & Schilling, K. Developmental Maturation of the Cerebellar White Matter-an Instructive Environment for Cerebellar Inhibitory Interneurons. Cerebellum 19, 286–308 (2020).

45. Zhan, Y., et al. Deficient neuron-microglia signaling results in impaired functional brain connectivity and social behavior. Nat Neurosci 17, 400–406 (2014).

46. Hoshiko, M., Arnoux, I., Avignone, E., Yamamoto, N. & Audinat, E. Deficiency of the microglial receptor CX3CR1 impairs postnatal functional development of thalamocortical synapses in the barrel cortex. J Neurosci 32, 15106–15111 (2012).

47. Zhang, Y., et al. An RNA-sequencing transcriptome and splicing database of glia, neurons, and vascular cells of the cerebral cortex. J Neurosci 34, 11929–11947 (2014).

48. Cunningham, C.L., Martinez-Cerdeno, V. & Noctor, S.C. Microglia regulate the number of neural precursor cells in the developing cerebral cortex. J Neurosci 33, 4216–4233 (2013).

49. Anderson, S.R., et al. Complement Targets Newborn Retinal Ganglion Cells for Phagocytic Elimination by Microglia. J Neurosci 39, 2025–2040 (2019).

50. Fünfschilling, U., et al. Glycolytic oligodendrocytes maintain myelin and long-term axonal integrity. Nature 485, 517–521 (2012).

51. Lee, Y., et al. Oligodendroglia metabolically support axons and contribute to neurodegeneration. Nature 487, 443–448 (2012).

52. Tang, G., et al. Loss of mTOR-dependent macroautophagy causes autistic-like synaptic pruning deficits. Neuron 83, 1131–1143 (2014).

53. Pei, J., et al. CX3CR1 mediates motor dysfunction in mice through 5-HTR2a. Behavioural brain research 461, 114837 (2024).

54. Pogarell, O., et al. EEG coherence reflects regional corpus callosum area in Alzheimer’s disease. J Neurol Neurosurg Psychiatry 76, 109–111 (2005).

55. Wegiel, J., et al. Brain-region-specific alterations of the trajectories of neuronal volume growth throughout the lifespan in autism. Acta Neuropathol Commun 2, 28 (2014).

56. Varghese, M., et al. Autism spectrum disorder: neuropathology and animal models. Acta neuropathologica 134, 537–566 (2017).

57. Billiards, S.S., et al. Development of microglia in the cerebral white matter of the human fetus and infant. J Comp Neurol 497, 199–208 (2006).

58. Verney, C., Monier, A., Fallet-Bianco, C. & Gressens, P. Early microglial colonization of the human forebrain and possible involvement in periventricular white-matter injury of preterm infants. J Anat 217, 436–448 (2010).

59. Craig, A., et al. Quantitative analysis of perinatal rodent oligodendrocyte lineage progression and its correlation with human. Exp Neurol 181, 231–240 (2003).

60. Oosterhof, N., et al. Homozygous Mutations in CSF1R Cause a Pediatric-Onset Leukoencephalopathy and Can Result in Congenital Absence of Microglia. American journal of human genetics 104, 936–947 (2019).

61. Rademakers, R., et al. Mutations in the colony stimulating factor 1 receptor (CSF1R) gene cause hereditary diffuse leukoencephalopathy with spheroids. Nat Genet 44, 200–205 (2011).

62. Nicholson, A.M., et al. CSF1R mutations link POLD and HDLS as a single disease entity. Neurology 80, 1033–1040 (2013).

63. Tada, M., et al. Characteristic microglial features in patients with hereditary diffuse leukoencephalopathy with spheroids. Ann Neurol 80, 554–565 (2016).

64. van der Knaap, M.S. & Bugiani, M. Leukodystrophies: a proposed classification system based on pathological changes and pathogenetic mechanisms. Acta neuropathologica 134, 351–382 (2017).

65. Bianchin, M.M., Martin, K.C., de Souza, A.C., de Oliveira, M.A. & Rieder, C.R. Nasu-Hakola disease and primary microglial dysfunction. Nat Rev Neurol 6, 2 p following 523 (2010).

66. Jay, T.R., von Saucken, V.E. & Landreth, G.E. TREM2 in Neurodegenerative Diseases. Molecular neurodegeneration 12, 56 (2017).

67. Wang, Y., et al. TREM2-dependent microglial function is essential for remyelination and subsequent neuroprotection. Glia 71, 1247–1258 (2023).

68. Olah, M., et al. Identification of a microglia phenotype supportive of remyelination. Glia 60, 306–321 (2012).

69. Yamanaka, K., et al. Deletion of Nox4 enhances remyelination following cuprizone-induced demyelination by increasing phagocytic capacity of microglia and macrophages in mice. Glia 71, 541–559 (2023).

70. Lampron, A., et al. Inefficient clearance of myelin debris by microglia impairs remyelinating processes. The Journal of experimental medicine 212, 481–495 (2015).

71. Gao, R., et al. Myelin debris phagocytosis in demyelinating disease. Glia (2024).

72. Deczkowska, A., et al. Disease-Associated Microglia: A Universal Immune Sensor of Neurodegeneration. Cell 173, 1073–1081 (2018).

73. Hammond, T.R., et al. Single-Cell RNA Sequencing of Microglia throughout the Mouse Lifespan and in the Injured Brain Reveals Complex Cell-State Changes. Immunity 50, 253–271 e256 (2019).

74. Li, Q., et al. Developmental Heterogeneity of Microglia and Brain Myeloid Cells Revealed by Deep Single-Cell RNA Sequencing. Neuron 101, 207–223 e210 (2019).

75. Kaiser, T., et al. MyelTracer: A Semi-Automated Software for Myelin g-Ratio Quantification. eNeuro 8(2021).

